# Subsurface carbon monoxide oxidation capacity revealed through genome-resolved metagenomics of a carboxydotroph

**DOI:** 10.1101/2020.06.25.170886

**Authors:** Andre Mu, Brian C. Thomas, Jillian F. Banfield, John W. Moreau

## Abstract

Microbial communities play important roles in the biogeochemical cycling of carbon in the Earth’s deep subsurface. Previously, we demonstrated changes to the microbial community structure of a deep aquifer (1.4 km) receiving 150 tons of injected supercritical CO_2_ (scCO_2_) in a geosequestration experiment. The observed changes support a key role in the aquifer microbiome for the thermophilic CO-utilising anaerobe *Carboxydocella,* which decreased in relative abundance post-scCO_2_ injection. Here, we present results from more detailed metagenomic profiling of this experiment, with genome resolution of the native carboxydotrophic *Carboxydocella.* We demonstrate a switch in CO-oxidation potential by *Carboxydocella* through analysis of its carbon monoxide dehydrogenase (CODH) gene before and after the geosequestration experiment. We discuss the potential impacts of scCO_2_ on subsurface flow of carbon and electrons from oxidation of the metabolic intermediate carbon monoxide (CO).

**Originality-Significance Statement:** The research conducted in this study is associated with one of the world’s largest demonstrations of carbon geosequestration – The Cooperative Research Centre for Greenhouse Gas Technologies Otway Project (Victoria, Australia). Our results expand the ecology of CO-utilising microbes to include the terrestrial deep subsurface through genome-resolved metagenomics of an aquifer-native carboxydotroph.

## Introduction

Carbon monoxide (CO) provides a carbon and/or energy source for a wide range of anaerobic microorganisms (Oelgeschläger and Rother, 2008) as its oxidation can be coupled to multiple terminal electron accepting processes (TEAPs), e.g., iron(III)-reduction (Slobodkin *et al.*, 1999), sulfate-reduction (Parshina *et al.*, 2005, 2010) hydrogenogenesis, acetogenesis or methanogenesis (Diender *et al.*, 2015). The electrons produced from CO oxidation can be coupled to the reduction of H_2_O and metals (Techtmann *et al.*, 2009). The substrate versatility of CO results from the low redox potential (E°; −524 to −558 mV) of the half-reaction for carbon dioxide (CO_2_) reduction to CO (Ragsdale, 2004; Techtmann *et al.*, 2009). This potential is lower than that of the H^+^/H_2_ redox couple, meaning that CO can replace H_2_ as an electron donor for microorganisms carrying genes encoding for carbon monoxide dehydrogenase (CODH). The CODH gene cluster in the genus, *Carboxydocella*, consists of 13 genes encoding for: redox- and CO-sensitive transcriptional regulator (*cooA*), [Ni, Fe]-CODH (*cooS*), [Ni, Fe]-CODH accessory nickel-insertion protein (*cooC*), CO-induced hydrogenase membrane anchor (*cooM*), energy-converging hydrogenase (*cooKLXUH*), hydrogenase maturation protein (*hypA)* 4Fe-4S di-cluster domain containing electron transfer protein (*cooF*), energy-converging hydrogenase catalytic subunit (*cooH*), and the upstream enigmatic *cooSC* operon of unknown function (Toshchakov *et al.*, 2018). The upstream *cooS* gene is notably missing in the closely related *Carboxydothermus hydrogenoformans* (Toshchakov *et al.*, 2018). The catalytic subunit of CODH is composed of the CooS protein, while the CooF electrontransfer protein is known to associate with redox reactions such as the production of H_2_ or reduced metals (Soboh *et al.*, 2002). Bi-functional CODH enzyme complexes contain domains that catalyse the oxidation of CO (CODH), and formation of acetyl-CoA (Acetyl-CoA synthase; ACS); Ragsdale and Kumar, 1996). Biochemically, the electrons produced by CODH flow on to the ACS module for downstream production of acetyl-CoA. Bi-functional CODH/ACS complexes are therefore fundamental to acetoclastic methanogens and acetogenesis in prokaryotes using the Wood-Ljungdahl pathway to form acetyl-CoA from either CO_2_ or CO (Ragsdale, 2004). The CODH/ACS complex is also responsible for the majority of autotrophic utilization of CO (Techtmann *et al.*, 2009). An increase in CO_2_ partial pressure may therefore inhibit microbial carboxydotrophy, resulting in increasing and decreasing levels of CO and H_2_, respectively. Such changes potentially hold consequences for both carbon and electron flow through various microbial metabolisms (Ragsdale, 2004; Techtmann *et al.*, 2009).

Using phenotypic and physiological assays, Sokolova et al. (2002) demonstrated that a 40% decrease in CO in cultures of the anaerobic Firmicute, *Carboxydocella,* lead to a 30% increase in the gas phase concentrations of H_2_ and CO_2_ (Sokolova *et al.*, 2002). Furthermore, pure cultures of *Desulfotomaculum kuznetsovii* and *D. thermobenzoicum* subsp. *thermosynthrophicum* were shown to grow on CO as the sole electron donor, and at concentrations as high as 50-70%, in the presence of hydrogen/CO_2_. However, co-culture of *Desulfotomaculum kuznetsovii* and *D. thermobenzoicum* subsp. *thermosynthrophicum* with a carboxydotroph (i.e, *Carboxydothermus hydrogenoformans)* supported sulfate-reducing bacteria (SRB) growth in 100% CO, with CO oxidation coupled to SO_4_^2-^ reduction and acetogenesis (pCO = 120 kPa) (Parshina *et al.*, 2005).

Injection of massive volumes of supercritical CO_2_ (scCO_2_) into deep aquifers forms a principal current strategy for geological carbon storage (Benson and Surles, 2006; Gibbins and Chalmers, 2008). These scCO_2_ “plumes” will enrich groundwaters locally with respect to dissolved CO_2_, with implications for both microbial metabolic activity and water-rock chemical reactions (Phillips *et al.*, 2012). Understanding the *in situ* impacts of increased CO_2_ on microbial CODH activity will yield insights into the long-term and large-scale potential responses of the subsurface microbial biosphere to geological CO_2_ sequestration.

Here we report results from a field-scale geological scCO_2_ injection project in the Paaratte Formation of the Otway Basin (1.4 km below ground, Southeastern Australia) that revealed a steep decline in the relative abundance of a dominant native carboxydotroph representing >96% of the aquifer microbial community before injection (Mu *et al.*, 2014). Previous studies of subsurface microbial responses to quickly-elevated CO_2_ levels have inferred their findings on the basis of 16S rRNA gene data (Bordenave *et al.*, 2013; Lavalleur and Colwell, 2013; Mu *et al.*, 2014). Here we used time-series relative abundance *in situ* metagenomic sampling to resolve a nearcomplete genome from the carboxydotrophic genus *Carboxydocella* and established the microbial host of the CODH gene cluster.

## 2. Results

### 2.1. Metagenomic analysis of the Paaratte Formation

A total of 13 samples collected from the Paaratte Formation over the course of a field-scale demonstration of supercritical carbon dioxide (scCO_2_) geosequestration were processed for shotgun metagenomics (Table 1) with multiple displacement amplification (MDA). Over five hundred tons of ground water were produced during the pre-scCO_2_ injection phase, followed by the injection of 150 tons of scCO_2_. Ten samples were collected during the pre-CO_2_ injection phase, two samples during the post-CO_2_ injection phase, and one sample from the coremud. The mean number of reads that passed quality control during the pre-scCO_2_ injection phase was 600,766, while for the post-scCO_2_ injection phase was 281,383. Mean GC content throughout the geosequestration experiment was 53%. The number of reads with identified functional categories was 76,662. The metagenome of the pre-CO_2_ injection phase sample PF13 contained 1,034,457 reads post-QC. Anaerobic CODH (represented by *cooS)* is absent from the two post-CO_2_ injection phase samples (Figure 1A), while aerobic CODH (represented by *coxL*) on the other hand was present at low levels during the pre-CO_2_ injection phase, but proliferated by ≥2.7 fold in sample PF145, which is during the late phases of post-CO_2_ injection (Figure 1B).

**Figure 1.**
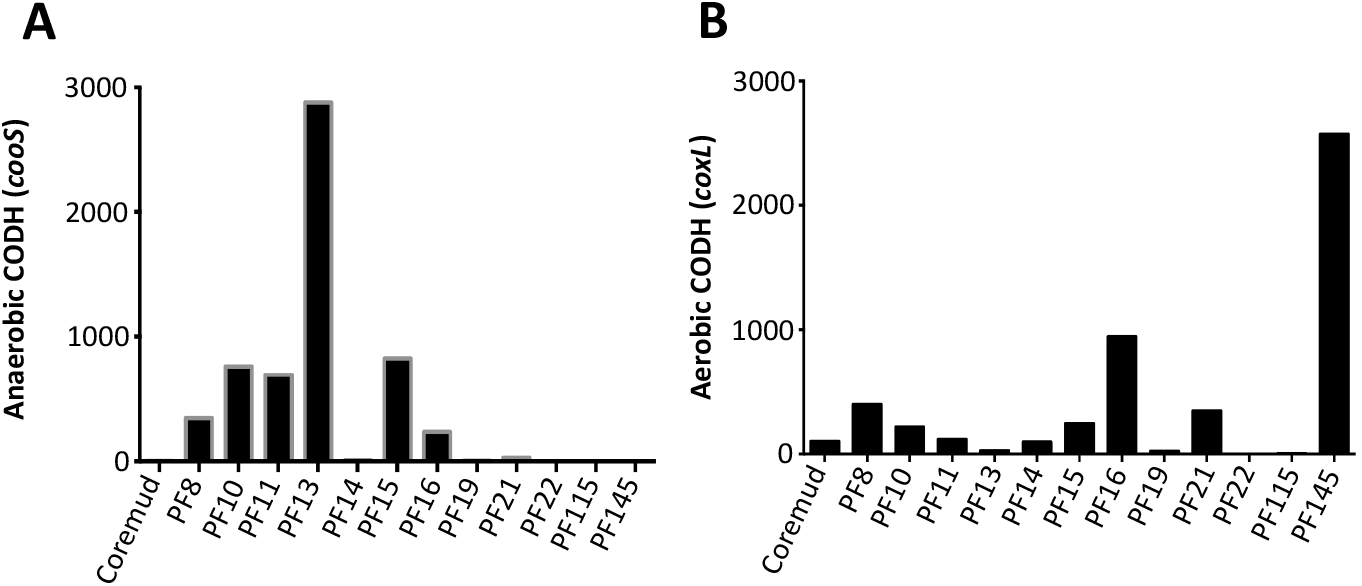
Abundances anaerobic and aerobic CODH genes throughout the geosequestration project. **(A)** An abundance profile of sequence data from the Paaratte Formation mapped to CooS. Coos is the catalytic subunit associated with the anaerobic carbon monoxide dehydrogenase eznyme. **(B)** An abundance profile of sequence data from the Paaratte Formation mapped to CoxL. CoxL is the large subunit associated with the aerobic carbon monoxide dehydrogenase eznyme.

**Table 1.**
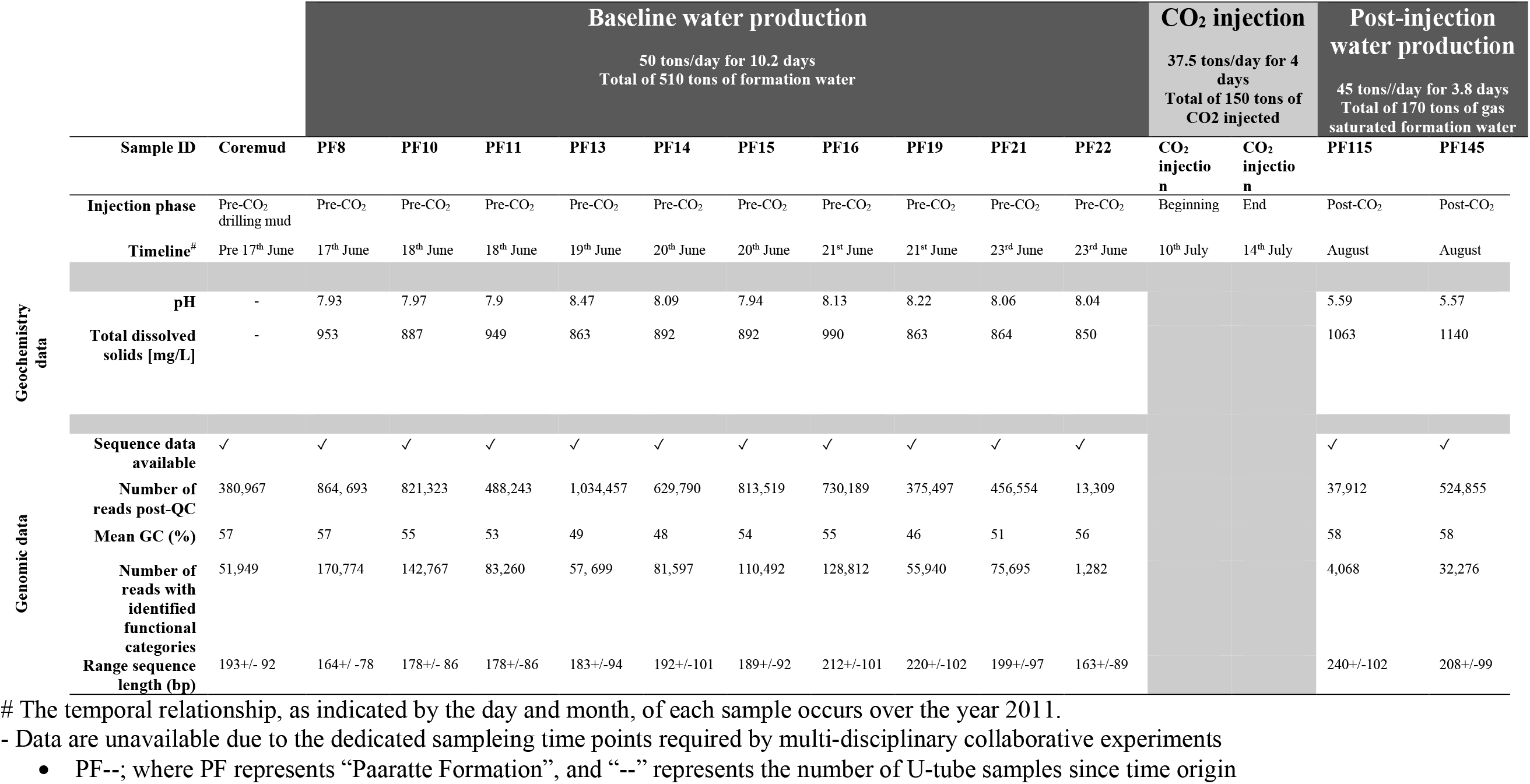
Summary of the thirteen samples analysed in this study

### 2.2. Assembly of metagenomic sequence data

Biodiversity analyses from our previous study (Mu *et al.*, 2014) informed the targeted selection of four samples from the pre-CO_2_ injection phase (samples PF10, 11, 13, and 15) for genome-resolved metagenomic analyses. Metagenomic assembly of these samples generated a total of 10,492 contigs with an average of 57.83% GC content and 31,465 predicted coding sequences (Table 2).

**Table 2.** Genomic features of metagenomic bins using pre-CO_2_ injection phase samples

| Samples | Completeness | Size | %GC | Average coverage | Number of contigs | Number of genes |
| --- | --- | --- | --- | --- | --- | --- |
| PF10 | **\*RP:** 47<br>**\*\*bSCG:** 47 | 10.29 Mbp | 60.4 | 16.1 | 4326 | 13,351 |
| PF11 | **RP:** 41<br>**bSCG:** 42 | 5.37 Mbp | 58.01 | 13.94 | 2582 | 7348 |
| PF13 | **RP:** 47<br>**bSCG:** 50 | 389.27 Kbp | 53.69 | 101.69 | 183 | 509 |
| PF15 | **RP:** 36<br>**bSCG:** 33 | 7.72 Mbp | 59.2 | 36.58 | 3401 | 10,257 |
| *Clostridiales* bins |  |  |  |  |  |  |
| PF10 | **RP:** 44<br>**bSCG:** 48 | 2.33 Mbp | 49.48 | 45.56 | 195 | 2521 |
| PF11 | **RP:** 42<br>**bSCG:** 46 | 2.13 Mbp | 49.49 | 37.05 | 216 | 2357 |
| PF13 | **RP:** 47<br>**bSCG:** 50 | 2.43 Mbp | 49.07 | 226.19 | 170 | 2659 |
| PF15 | **RP:** 35<br>**bSCG:** 38 | 1.95 Mbp | 50.05 | 59.54 | 427 | 2318 |

\*RP = Ribosomal proteins
\*\*bSCG = bacterial single copy genes

### 2.3 Metagenomic reconstruction of aquifer-native carboxydotroph

Metagenomic bins were created for a carboxydotroph *Carboxydocella* using ggKbase from samples PF10, PF11, PF13, and PF15, based on the following bin phylogenetic profile; Bacteria > *Firmicutes* > *Clostridia* > *Clostridiales;* and GC contents (%) of 43.56 – 51.99 for PF10, 40.7 – 53.98 for PF11, 43.13 – 55.0 for PF13, and 39.43 – 51.5 for PF15. The metagenomic bin for *Carboxydocella* from PF13 had the most bacterial single copy genes (bSCGs) (50 of 51 expected bSCGs). This sample contained highest relative abundances of *Carboxydocella*, as determined by amplicon 16S rRNA gene profiling. Samples PF10, PF11, and PF15 returned *Carboxydocella* genomes with 48, 46 and 38 of each of the 51 expected bSCGs, respectively (Table 1). Notably, the gene encoding for the catalytic subunit of CODH enzyme (*cooS)* was present in the genomes from all four metagenomic datasets (Figure 2B); also present were genes associated with *Flagella related proteins*.

**Figure 2.**
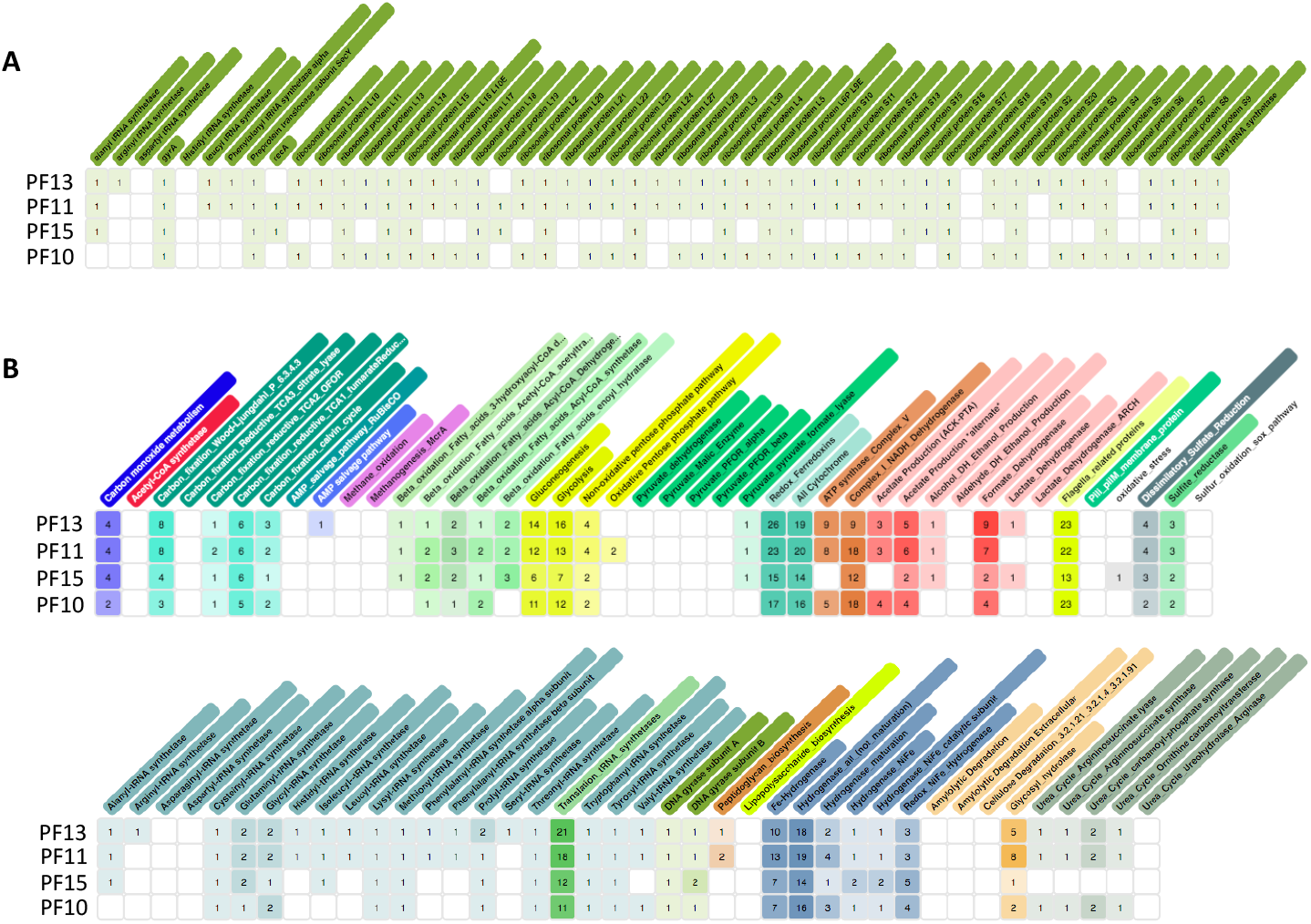
Genomic characterization of genome-resolved aquifer-native carboxydotroph metagenome. **(A)** Bacterial single copy gene markers for metagenomic bins representing the *Carboxydocella* operational taxonomical unit. Bacterial single copy genes are used as genetic biomarkers to validate the genome is derived from a single community isolate, and as a proxy for genome completeness. Contigs were clustered together using the ggKbase analysis tool. **(B)** Genomic features of aquifer-native carboxydotrophic metagenomic bins. Each row is representative of a *Carboxydocella* genome from a single time point of the geosequestration project. Presence of a genomic feature is indicated by a coloured cell, and its abundance by the numerical value. Row 1 = Sample PF13; Row 2 = Sample PF11; Row 3 = Sample PF15; Row 4 = Sample PF10.

The manually curated genome-resolved metagenome of the aquifer-native carboxydotroph is 2,544,661 bp in length, consisting of six rRNA genes, 64 tRNAs, 2584 coding sequences (CDS), and has a GC content of 49.2% GC. These genomic characteristics are similar to those of the two reference *Carboxydocella* strains. The genome is classified as near-complete (94.6% complete) when assessed using *C. thermautotrophica* strain 019 as the reference genome (Table 3). BLASTP and phylogeny-based analyses revealed high similarity between aquifer-native carboxydotroph and the two reference strains (*C. thermautotrophica* strain 019, accession number: CP028491; and strain 41, accession number: CP028514). The genome has the complete gene cluster determining hydrogenic carboxydotrophy: *cooA*-transcriptional regulator; *cooS* – Ni,Fe-CODH gene; *cooC* – Ni,Fe-CODH accessory nickel-insertion protein gene; *cooMKLXUH* – energy-converting hydrogenase gene; *hypA*-hydrogenase maturation protein gene; *cooF* ferredoxin-like pretien gene; and *cooH* – ECH catalytic subunit gene. The gene cluster also includes an upstream enigmatic *cooSC* operon, in which the function in *Carboxydocella* is unknown (Figure 2D). The reconstructed aquifer-native *Carboxydocella* genome is available under the Europen Nucleotide Archive (ENA) study accession number: PRJEB37484.

**Table 3.** Genomic features of the aquifer-native *Carboxydocella* compared to *C. thermautotrophica* reference strains

|  | Genome length (bp) | %GC | CDS | rRNA | tRNA | Number of contigs | Ns | Min contig | Max contig | N50 |
| --- | --- | --- | --- | --- | --- | --- | --- | --- | --- | --- |
| **PF-AMU\*** | 2,544,661 | 49.2 | 2584 | 6 | 64 | 55 | 15 | 1369 bp | 252,013 bp | 89,432 bp |
| ***C. thermautotrophica* 019** | 2,676,584 | 49.1 | 2689 | 15 | 71 |  |  |  |  |  |
| ***C. thermautotrophica* 41** | 2,690,058 | 49.1 | 2697 | 15 | 73 |  |  |  |  |  |

\* Near-complete genome-resolved metagenome of aquifer native *Carboxydocella*

## 3. Discussion

Analysis of 16S rRNA gene data revealed a proliferation of *Comamonadaceae* and *Sphingomonadaceae*, and a decline of *Carboxydocella*, following the injection of scCO_2_ to the Paaratte Formation (Mu *et al.*, 2014). Temporal shifts in phylogeny also related to changes in metabolic potential, and provided insights into likely effects of scCO_2_ on CO metabolism by *Carboxydocella* (Mu *et al.*, 2014). *Carboxydocella* is a Gram positive *Firmicute* that has been phenotypically (including biochemically) described by Sokolova and colleagues (2012) to be motile (lateral flagella), and capable of oxidising CO as its sole carbon source. However, there are few sequenced reference strains available in public databases for attributing the molecular mechanisms behind the oxidation of CO in terrestrial subsurface environments (Doukov *et al.*, 2002; Matson *et al.*, 2011).

Metagenomic “binning” of sequence data based on organism-specific characteristics (e.g., GC content, phylogeny, coverage, and bacterial single copy genes), and manual curation of assembly data produced high-quality genomes for an aquifer-native carboxydotroph; this genome represents a near-complete genome at 94.6% complete compared to *C. thermautotrophica* strain 019, and at 49.2 percentage GC (Figure 3A, 3B, 3C). Despite our necessary use of MDA to prepare genomic samples for sequencing, which has inherent biases to GC% content and taxonomy, the metagenomic bins and reconstructed carboxydotroph genome demonstrated comparable genomic characteristics to reference isolates (Toshchakov *et al.*, 2018). Reconstruction of the genome for aquifer-native carboxydotrophs revealed the presence of anaerobic Ni-dependent CODH CooS (*cooS*) belonging to *Carboxydocella*, supporting the capacity of this organism for CO oxidation (Figure 3D). Consistent with expectations based on genomes of related bacteria, the aquifer-native carboxydotroph genome encodes flagella related proteins and lacks lipopolysaccharide biosynthetic genes (Figure 2B); these key genomic features support the phenotypic characterization of *Carboxydocella* being motile via lateral flagella, and having a cell wall structure consistent of a Gram positive microorganism (Sokolova *et al.*, 2002).

**Figure 3.**
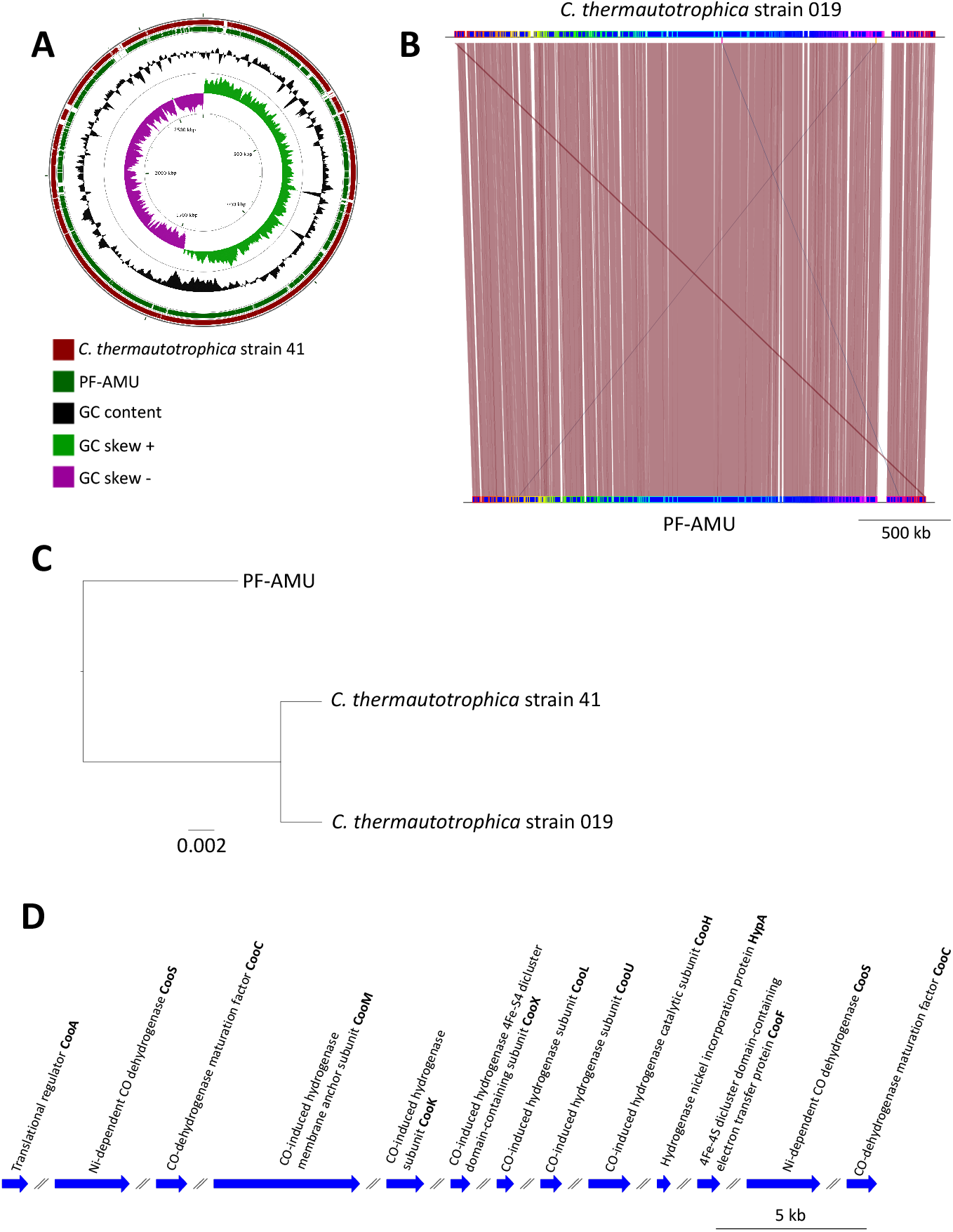
Comparative analysis of the aquifer-native carboxydotroph with representative carboxydotrophic microbial species. **(A)** CGView tools (BLASTP) analysis of *Carboxydocella thermautotrophica* strains 41 and the aquifer-native carboxydotroph with *C. thrermautotrophica* str 019 as the reference. **(B)** MauveProgressive aligner comparasion of aquifer-native carboxydotroph (PF-AMU) and *C. thrermautotrophica* str 019 genomes as visualized by genoPlotR. **(C)** Core SNP phylogenetic tree of *C. thermautotrophica* strains 41 and 019, and the aquifer-native carboxydotroph. Variant analysis was computed using Prokka for *C. thermautotrophica* strain 41 and the aquifer-native carboxydotroph against *C. thermautotrophica* strain 019 as the reference. **(D)** Schematic of the gene cluster determining hydrogenic carboxydotrophy in the aquifer-native *Carboxydocella* genome. The gene cluster is homologous to that of *C. thermautotrophica* strain 019 and 41.

Metagenomic data revealed that representative aerobic CODH, *coxL,* genes proliferated during the late phase of post-scCO_2_ injection by ≥2.7 fold (Figure 1B). The observed increase in aerobic CODH may have resulted from a decline in the activity of *Carboxydocella,* which cannot grow in the presence of large volumes of CO_2_ (Sokolova *et al.*, 2002). However, proliferation of *coxL,* and a decline in *cooS*, suggests a metabolic switch in CO-oxidation potential within the subsurface. CO-oxidation is likely maintained by microbes other than anaerobic carboxydotrophs, such as those using molybdenum containing CO oxidoreductase enzymes (Rajagopalan, 1984; King and Weber, 2007). This observed switch in CO-oxidation potential, rather than a complete loss of function within the aquifer community, implies that sulfate-reducing bacteria and methanogens would have sufficient H_2_ to sustain physiological function (Mu and Moreau, 2015). Furthermore, few aerobic carboxydotrophs have been documented in thermophilic environments (King and Weber, 2007); therefore, our findings extend the range of geological environments for aerobic CO-oxidisers.

This study offers insights for future *in vitro* studies of pure culture and mixed consortia responses to scCO_2_. Our results elucidate the roles of anaerobic and aerobic CODH in microbial community functionality in a system under scCO_2_ stress, and expand the ecology of CO-utilising microbes to include the terrestrial deep subsurface. Furthermore, the genome-resolved metagenome of a Paaratte Formation-derived *Carboxydocella* organism provides a greater molecular understanding of CO-oxidation within the deep subsurface.

## 4. Experimental procedures

### 4.1 Groundwater sampling and extraction of whole community genomic DNA

*In situ* groundwater samples were collected using the U-tube sampling system (Freifeld *et al.*, 2005; Freifeld, 2009) as part of the CO_2_CRC Otway Stage 2B field experiment (Paterson *et al.*, 2013). Details of groundwater sampling and extraction of whole community genomic DNA (gDNA) are given in Mu et al., (2014). The injection and sampling well was screened at approximately 1400 meters true vertical depth subsea (TVDSS) in a sandstone unit of the Paaratte Formation of the Otway Basin at latitude: 38° 31’ 44” and longitude: 142° 48’ 43”. A series of valve controls at surface were used to pass high-pressure nitrogen gas through the U-tube system and direct formation water to the surface for collection under under approximately 2000 psi. Each sample collected via the U-tube system was designated the nomenclature “PF---”; where “PF” represents *Paaratte Formation,* and “—” represents the number of U-tube samples since time of origin, which was taken as 17^th^ June 2011. Control samples included nucleic acid extracted from core mud, which is representative of drilling fluid used during the emplacement of injection and sampling wells, and samples from water tanks used to store formation water at the surface. Samples processed for metagenomic analysis are highlighted in Table 1.

### 4.2. Preparation and processing of genomic DNA

#### 4.2.1. Whole genome amplification

To increase the amount of high-quality gDNA to meet input requirements for illumina’s Nextera XT sample preparation kit, it was necessary to amplify extracted gDNA using MDA (Warnecke and Hess, 2009; Kawai *et al.*, 2014) with Qiagen’s REPLI-g UltraFast Mini Kit following manufacturer’s protocol. Genomic DNA from samples PF8, PF10, PF11, PF13, PF14, PF15, PF16, PF19, PF21, PF22, PF115, PF145, and coremud, were amplified using MDA.

#### 4.2.2. Genomic DNA preparation for metagenomic sequencing

Sample preparation and metagenomic sequencing were performed using the Illumina MiSeq located in the Department of Microbiology and Immunology at the Peter Doherty Institute for Infection and Immunity, University of Melbourne, Australia.

Multiple displacement amplified gDNA was quantitated using the Qubit dsDNA BR (Molecular probes^®^) assay system following manufacturer’s protocol. Samples of sufficient quality were processed using illumina’s Nextera XT sample preparation kit to generate Clean Amplified Nextera Tagment Amplicons following the manufacturer’s protocol. Clean Amplified Nextera Tagment Amplicon-DNA concentrations were checked using Qubit dsDNA High Sensitivity Kit, while DNA fragment sizes were validated and quantified using the Agilent 2100 Bioanalyzer and Agilent high sensitivity DNA kit. Table S1 highlights the dilution factors for each sample library to obtain correct concentrations for sequencing on the MiSeq Sequencer. Samples were diluted using Qiagen’s EB (elution buffer) in place of Tris-Cl 10 mM 0.1% Tween 20.

#### 4.2.3. Illumina MiSeq sequencing

Samples were processed for sequencing using the Illumina MiSeq reagent kit v2 (500 cycle) on the MiSeq machine following a modified manufacturer’s protocol. The following modifications were included: a 1% (v/v) spike-in ratio of PhiX, denatured DNA was diluted to a final concentration of 12 pM with pre-chilled HT1 buffer, and Tris-Cl 10 mM 0.1% Tween 20 was substituted with Qiagen’s EB solution to dilute sequencing libraries and PhiX throughout the protocol. Raw sequencing data is available through the Europen Nucleotide Archive (ENA) study accession number: PRJEB37484.

### 4.3. Metagenomic analysis

Pre-processing steps included the removal of artificial sequences generated by sequencing artifacts, and filtering sequence reads that mapped to the *Homo sapiens* genome from the library intended for further analysis using Bowtie (Langmead and Salzberg, 2012). The pre-processing step aimed to remove contaminating sequences derived from the handling and preparation of gDNA libraries. Sequences were trimmed to contain at most five bases below a Phred score of 15, which was considered to be the lowest quality score included as a high-quality base. Furthermore, the maximum allowed number of ambiguous base pairs per sequence read was five.

Illimuna sequence data were processed with Trimmomatic (version 0.33; (Bolger *et al.,* 2014) for quality control and to remove adaptor sequences, PhiX contamination, and trace contaminants from illumina preparation kits. Paired-end reads were merged and assembled using Iterative de Bruijn Graph De Novo Assembler for Uneven sequencing Depth (IDBA-UD; Peng *et al.*, 2012)) compiled for long reads (i.e., 351 bp). Scaffold headers in data files were amended to include read mapping coverage calculations from Bowtie2 (version 2.2.4; Langmead and Salzberg, 2012) using a ruby script (*add_read_count.rb*; source code available from https://github.com/bcthomas/misc_scripts/tree/master/add_read_count) written by University of California, Berkeley researchers. Prokaryotic Dynamic Programming Gene finding Algorithm (Prodigal v2.6.3; (Hyatt *et al.*, 2010) was utilised to predict genes and small RNAs from assembled sequence data greater than 1000 bp in length. Annotation of the predicted proteins were computed by doing similarity searches using *usearch* (v8.1.1861) (Edgar, 2010), and comparing each protein sequence against the following databases: Kyoto Encyclopedia of Genes and Genomes (KEGG; release 71.1 (Kanehisa and Goto, 2000), Consortium, 2009) and Universal Protein Resource (UniProt release 2014_08; Consortium, 2009) including Universal Protein Resource Reference Clusters (UniRef100; Consortium, 2009). Metagenomic data from samples PF10, PF11, PF13, and PF15, were processed for analysis using ggKbase (http://ggkbase.berkeley.edu/) to reconstruct the aquifer-native carboxydotroph. Key metabolic features were analysed against each genome across samples PF10, PF11, PF13, and PF15.

Computational requirements were provided by the Victorian Life Sciences Computation Initiative (Melbourne, Australia), and the University of California, Berkeley (California, United States). Manual curation of the metagenomic bin corresponding to the aquifer-native carboxydotroph was computed using Geneious (v 9.1.8; Biomatters, New Zealand). Prokka was used to annotate the aquifer-native carboxydotroph; annotations were guided by the reference *C. thermautotrophica* strain 019 GenBank file (Accession number: CP028491). Coordinates for the CODH gene cluster were identitifed and mapped using genoPlotR. CGViewtools and Mauve were also used to visualize genome comparisons between the aquifer-native carboxydotroph, *C. thermautotrophica* strains 019, and 41.

## Acknowledgements

The authors acknowledge funding to JWM from the Commonwealth of Australia and industry sponsors through the CO_2_CRC Program, and the Victorian Life Sciences Computation Initiative (grant number VR0032).

